# Systematic genome-guided discovery of antagonistic interactions between archaea and bacteria

**DOI:** 10.1101/2024.09.18.613068

**Authors:** Romain Strock, Valerie WC Soo, Antoine Hocher, Tobias Warnecke

## Abstract

The social life of archaea is poorly understood. In particular, even though competition and conflict are common themes in microbial communities, there is scant evidence documenting antagonistic interactions between archaea and their abundant prokaryotic brethren: bacteria. Do archaea specifically target bacteria for destruction? If so, what molecular weaponry do they use? Here, we present an approach to infer antagonistic interactions between archaea and bacteria from genome sequence. We show that a large and diverse set of archaea encode peptidoglycan hydrolases, enzymes that recognize and cleave a structure – peptidoglycan – that is a ubiquitous component of bacterial cell walls but absent from archaea. We predict the bacterial targets of archaeal peptidoglycan hydrolases using a structural homology approach and demonstrate that the predicted target bacteria tend to inhabit a similar niche to the archaeal producer, indicative of ecologically relevant interactions. Using a heterologous expression system, we demonstrate that two peptidoglycan hydrolases from the halophilic archaeaon *Halogranum salarium* B-1 kill the halophilic bacterium *Halalkalibacterium halodurans,* a predicted target, and do so in a manner consistent with peptidoglycan hydrolase activity. Our results suggest that, even though the tools and rules of engagement remain largely unknown, archaeal-bacterial conflicts are likely common, and we present a roadmap for the discovery of additional antagonistic interactions between these two domains of life. Our work has implications for understanding mixed microbial communities that include archaea and suggests that archaea might represent a large untapped reservoir of novel antibacterials.

## INTRODUCTION

Archaea are best known as extremophiles, thriving in environments where few other organisms survive. Such environments – from hot springs to salt lakes to acid mine drainage - are often dominated or even exclusively inhabited by archaea (Distaso et al. 2020; Belilla et al. 2021; Sriaporn et al. 2023). Over the last 10-15 years, however, broader and more sensitive sampling has revealed that archaea are not limited to such extreme environments. On the contrary, they are found in diverse locations including the digestive tracts of cattle and termites, where they drive methane production, and various soil and marine ecosystems where they play key roles in global carbon and nitrogen cycles (Moissl-Eichinger et al. 2018). In all of these settings, archaea are surrounded – and typically outnumbered – by bacteria (Medina-Chávez and Travisano 2022).

How archaea interact – physically, ecologically, and molecularly - with bacterial members of their respective communities is poorly understood (Moissl-Eichinger *et al*. 2018). What we do know largely concerns commensal or symbiotic associations, where at least one member of the community benefits from the interaction and nobody is harmed. Notably, this includes syntrophic consortia of methane-oxidising archaea and sulphate-reducing bacteria in marine environments, where archaea consume end products of bacterial metabolism (e.g. acetate, hydrogen, or formate) that would otherwise inhibit bacterial growth (Knittel and Boetius 2009; Wegener et al. 2022). Syntrophic interactions have also been observed inside eukaryotic hosts, including the guts of mammals (Samuel and Gordon 2006; Wrede et al. 2012; Ruaud et al. 2020), and likely underpin some intimate physical associations between archaea and bacteria (Rudolph et al. 2001; Rudolph et al. 2004; Muller et al. 2010).

One might suspect, however, that things are not always harmonious. For most niches, conflict is probably more common than cooperation (Mitri and Foster 2013; Palmer and Foster 2022). Yet evidence for antagonistic interactions between archaea and bacteria (where one species harms the other, be that through competition, predation, or incidentally) is very limited. We are aware of only a handful of reports detailing antagonistic interactions, all involving supernatant from archaea grown in pure culture, which was found to inhibit the growth of (often ecologically unrelated) indicator bacteria (Atanasova et al. 2013; Megaw et al. 2019; Castro et al. 2021; Liang et al. 2023). None of these studies determined the mechanism of inhibition. Nor did they answer the question whether antibacterial activity had evolved as a specific adaptation or whether bacteria were simply innocent by-standers, killed in the crossfire of mechanisms evolved to target other archaea (or perhaps eukaryotes).

Does the dearth of reported cases of archaeal-bacterial conflict imply that archaea generally shun confrontation and that niche partitioning is largely frictionless and amicable? To us, this seems unlikely; archaea, after all, are not shy to defend their niche against other *archaea*, using a variety of proteins or small molecules whose precise nature and mechanisms of action remain largely unknown (Torreblanca et al. 1994; Meseguer et al. 1995; Shand and Leyva 2007; Atanasova et al. 2013; Besse et al. 2015).

How might one determine whether archaea antagonistically – and specifically – target bacteria? We reasoned that one might do so by looking, in archaeal genomes, for proteins that destructively target cellular structures that are unique to bacteria. One prominent such structure is peptidoglycan, a continuous mesh that envelops the cytoplasmic membrane of bacterial cells, composed of polysaccharide chains that are covalently crosslinked by peptide moieties. Some methanogenic archaea produce analogous extracellular structures of cross-linked glycans and peptides, but this pseudo-peptidoglycan (pseudomurein) exhibits different composition and linkage patterns, making it chemically distinct from bacterial peptidoglycan (Mukhopadhyay 2024). We therefore decided to survey archaeal genomes for the presence of enzymes that cleave bacterial peptidoglycan: peptidoglycan hydrolases (PGHs).

PGHs are ubiquitous in bacteria. They typically exhibit modular architectures (Fig. 1A), with one or more peptidoglycan binding domains, which confer target specificity, fused to a catalytic domain (i.e. an amidase, peptidase, or glycosidase), which cleaves specific bonds of the peptidoglycan molecule. Bacteria deploy PGHs to remodel their own cell wall during cell division (autolysins), but occasionally also as weapons (Uehara and Bernhardt 2011). In these instances, following secretion, the PGH no longer acts on the producer’s own peptidoglycan but diffuses to its target – often a closely related strain (Akesson et al. 2007; Azevedo et al. 2015) – where it cleaves peptidoglycan to compromise cell wall integrity. Examples of such weaponized PGHs include lysostaphin from *Staphylococcus simulans* bv. staphylolyticus (Fig. 1A) and zoocin A from *Streptococcus equi* subsp*. zooepidemicus* 4881, which act on a number of other Staphylococcus and Streptococcus strains, respectively (Wu et al. 2003; Akesson et al. 2007).

**Figure 1.**
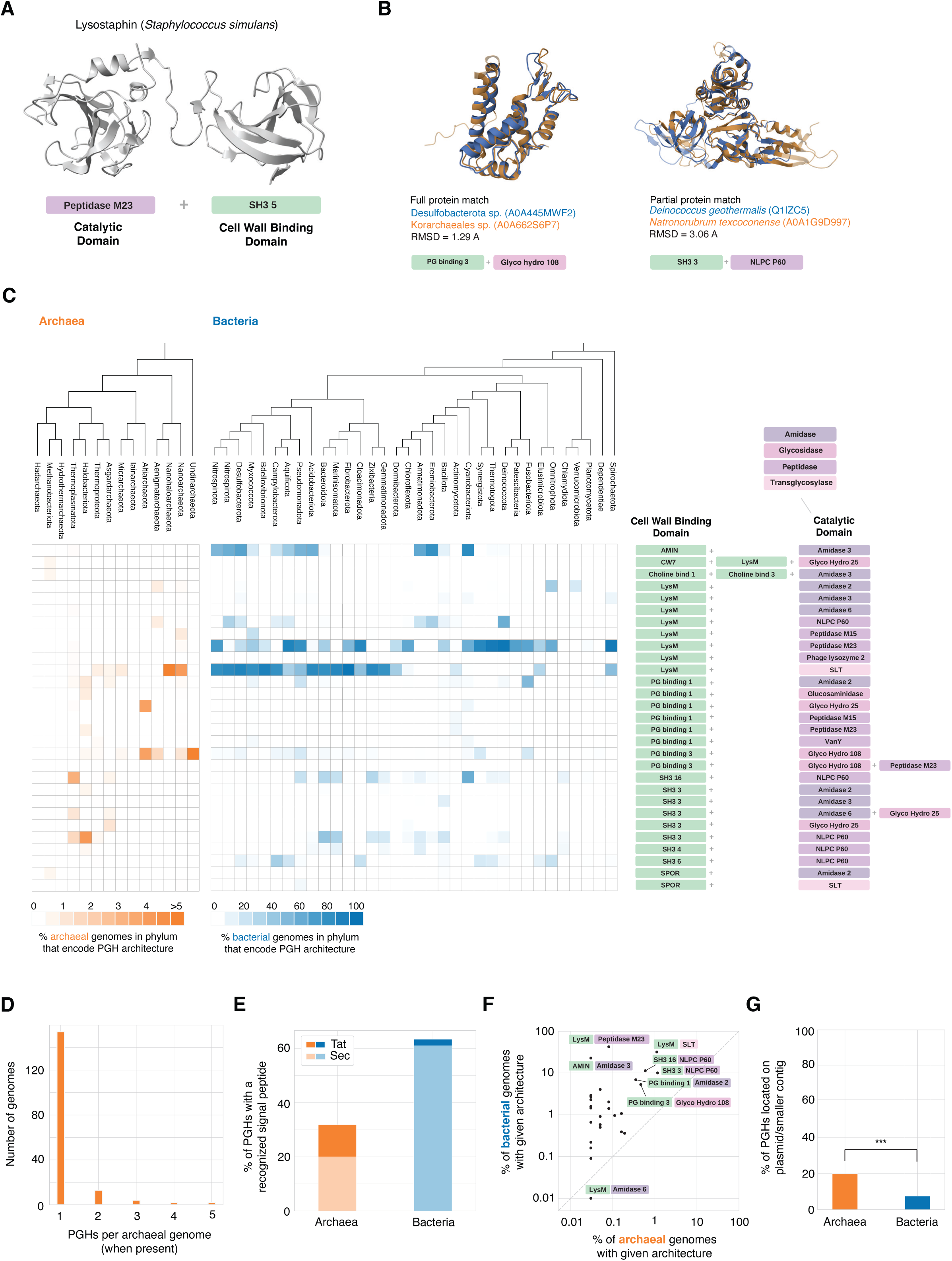
Peptidoglycan hydrolases in archaea. **A.** Structure of the peptidoglycan hydrolase (PGH) lysostaphin from *Staphylococcus simulans*, highlighting its modular domain architecture. **B.** Examples of structural homology between bacterial and archaeal homologs, covering either the full (left) or parts of (right) the protein query. **C.** Distribution of PGH homologs across archaea and bacteria, summarizing the relative abundance of different domain combinations at the phylum level. **D.** Number of archaeal genomes in the database that encode one or more PGHs. **E.** Proportion of PGHs with a known signal peptide, split according to the predicted secretion system (Tat/Sec). **F.** Incidence of different PGH architectures in bacteria versus archaea. **G.** Proportion of PGH homologs located on a plasmid (for completely assembled genomes) or, for incompletely assembled genomes, contigs below the median size of contigs in the assembly. ***P=8.9*10^-7^, Fisher’s exact test.

We reasoned that, since archaea do not have peptidoglycan, the presence of peptidoglycan-cleaving proteins in archaeal genomes might suggest that these archaea interact with bacteria in an antagonistic manner. Indeed, one case of an archaeon-encoded PGH with antibacterial activity has previously been reported: tracing horizontal gene transfer of antibacterial genes across the tree of life, Metcalf and colleagues discovered a GH25 muramidase (UniProt ID: B5ID12) in the genome of *Aciduliprofundum booneii* (Metcalf et al. 2014). A purified version of this PGH was shown to kill a range of gram-positive bacteria in the family Bacillaceae, from which the enzyme was likely acquired. Below, we show that the case of *A. booneii* B5ID12 is not a collector’s item and that PGHs with bactericidal activity are found in the genomes of diverse archaea, with implications for our understanding of how archaea act within and shape polymicrobial ecosystems.

## RESULTS

### Archaeal genomes harbour diverse peptidoglycan hydrolases

We assembled and surveyed a phylogenetically balanced database of prokaryotic genomes (3,706 archaea; 50,640 bacteria, Table S1) for proteins with homology to known PGHs (see Methods). As expected, we find recognizable PGHs in a large proportion of bacterial genomes (N=42,612 — 84%). Surprisingly, however, we also find PGH homologs in nearly 5% (N=175/3,706) of archaeal genomes, including representatives from all archaeal superphyla and spanning a broad range of PGH architectures (Fig. 1C; Table S2). PGH homologs are particularly prominent (and diverse) amongst Nanoarchaeota, which are thought to lead mostly symbiotic or parasitic lives (Dombrowski et al. 2019), Thermoplasmatota, and Halobacteriota. While not the rule, it is not uncommon for a single archaeal genome to encode more than one PGH (12%, N=21/175, Fig. 1D; Table S2).

For archaeal PGHs to act on bacterial peptidoglycan, they need to be secreted. We find that 32% (N=67/210) of archaeal PGH homologs encode a known signal peptide (65% Sec; 35% Tat; Fig. 1E; Table S2), consistent with deployment outside the cell.

In some instances, the binding domain of a PGH may interact with substrates other than peptidoglycan. For example, the LysM domain can, at least on occasion, bind chitin as well as peptidoglycan (Visweswaran et al, 2012), and some archaea secrete glycosyl hydrolases to digest chitin extracellularly as an alternative carbon source (Cono et al. 2020). In many cases, however, structural homology (see Fig. 1B for examples), conservation of known catalytic residues, and the presence of both a peptidoglycan binding domain *and* a catalytic domain involved in peptidoglycan cleavage suggest peptidoglycan as the conserved substrate.

To investigate the functionality of archaeal PGH homologs in greater detail, we decided to focus on archaeal homologs of zoocin A, one of the best-described weaponized PGHs in bacteria (Lai et al. 2002; Heath et al. 2004; Akesson et al. 2007; Simmonds et al. 2009). We detect homology against the peptidase M23 domain of zoocin A in ten archaeal genomes in our database, including in five genomes belonging to the order Halobacteriales (Table S2). The cell wall binding domain of *S. equi* zoocin A (zoocin A target recognition domain, PF16775) is not found in archaea. Constructing a pan-prokaryotic phylogenetic tree of the M23 domain on its own, we find that the homologous domains found in Halobacteriales form two monophyletic groups, each branching with different bacteria from the phylum Bacillota (Fig. 2A). In these archaea, the M23 domain lies N-terminal of two tandem peptidoglycan binding domains (PG binding 1 domain; PF01471). In the most closely related bacterial proteins, on the other hand, we find M23 C-terminal of a LysM domain. These observations suggest an evolutionary history marked by domain shuffling, previously proposed as a hallmark of bacterial PGHs (Diaz et al. 1991). A phylogenetic tree built from the PG binding 1 domain supports this view, highlighting an entirely different set of bacteria, some of which exhibit the same domain composition found in archaea (Fig. 2B).

**Figure 2.**
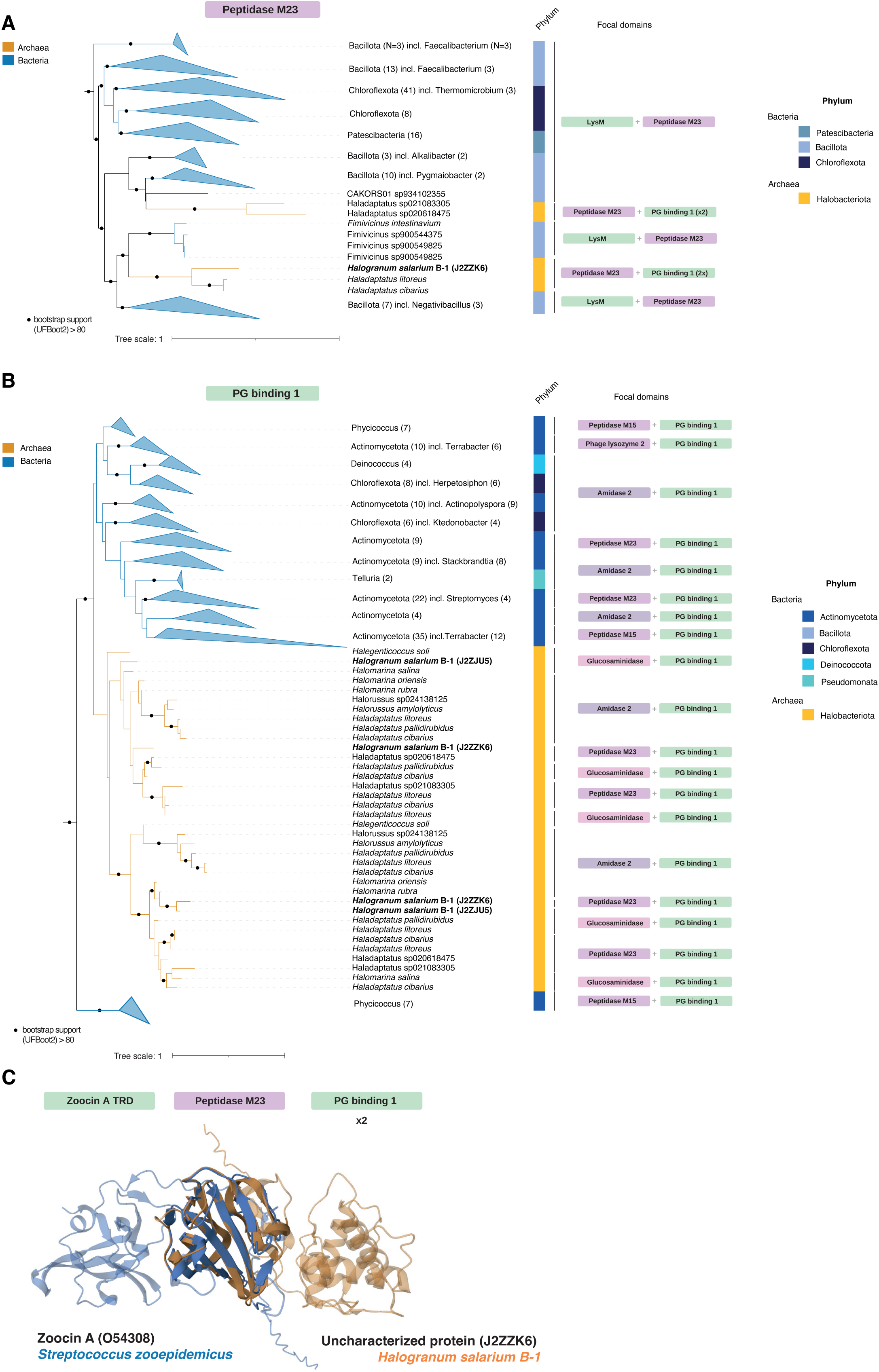
Chimeric zoocin A homologs in archaea. Protein domain-level phylogenetic trees highlighting evolutionary relationships of **(A)** peptidase M23 (Pfam ID: PF0155) and **(B)** PG binding 1 domains (Pfam ID: PF01471). The trees shown do not include all homologs present in archaea/bacteria but are subtrees that include all archaeal homologs from the Halobacteriota phylum and their immediate bacterial context. Domains within each protein are treated individually; hence proteins with two or more instances of the focal domain appear more than once in the tree. **C.** Structural overlay of zoocin A from *Streptococcus zooepidemicus* (UniProt ID: O54308) and a candidate PGH from *Halogranum salarium* B-1 (UniProt ID: J2ZZK6), highlighting homologous M23 domains but divergent cell wall binding domains. TRD: target recognition domain. PG: peptidoglycan. See Methods for details on how trees were constructed. Note that domain composition describes the presence of a given domain but not its number. Most proteins here carry more than one copy of a particular cell wall binding domain, including J2ZZK6, which has two tandem copies of the PG binding 1 domain.

The closest bacterial sister clade to the archaeal PG binding 1 domains contains a phylogenetic jumble of bacteria – including Actinomycetota, Chloroflexota, and Deinococcota – and a variety of domain architectures (Fig. 2B) suggesting that horizontal transfer and domain shuffling have also been happening amongst bacteria.

Although we find the M23 domain in several Halobacteriales genomes, the majority of Halobacteriales genomes in our database do not encode this domain (Fig. S1). This patchy phylogenetic distribution, where homologs are present in some closely related genomes but absent from others, is not dissimilar from that of weaponized PGHs, and bacteriocins more generally, in bacteria (Azevedo et al. 2015), where it has been interpreted as consistent with evolution under an arms-race regime: PGH utility changes rapidly as targets adapt and new targets emerge, driving frequent gain and loss.

Transient utility of PGHs and their exchange via HGT is further supported by the fact that a significant fraction is found on plasmids versus main chromosomes (4/18 in archaea compared to 102/13,786 in bacteria; Fisher’s exact test p-value < 1e-5; Fig. 1G), or – for genomes that are incompletely assembled – enriched on smaller contigs, which are *a priori* more likely to correspond to plasmids or secondary chromosomes (26/133 in archaea vs 8,292/97,918 in bacteria; Fisher’s exact test p-value < 1e-4; Fig. 1G, see Methods).

Taken together, a patchy phylogenetic distribution (Fig. S1), biased localization on plasmids (Fig. 1G), capacity for secretion (Fig. 1E), and high levels of structural conservation (e.g. Fig. 1B) are consistent with the idea that archaeal PGHs might be used in conflicts with bacteria.

### A peptidoglycan hydrolase from Halogranum salarium B-1 kills a halophilic bacterium

Next, we sought to establish whether archaeal PGHs do indeed have antibacterial activity and, if so, which bacteria are targeted. To this end, we decided to focus on one of the chimeric zoocin A homologs we identified, from the halophilic archaeon *Halogranum salarium* B-1 (Uniprot ID: J2ZZK6, Fig. 2C), which can be cultured with relative ease in the laboratory.

With an archaeal PGH homolog in hand, how can we identify the likely bacterial target(s)? We reasoned that, as the peptidoglycan-binding (PGB) domain of the PGH confers target specificity (Baba and Schneewind 1996; Mitkowski et al. 2019), we might home in on putative target bacteria by looking for homology between the PGB domain of a given archaeal PGH and PGB domains encoded in bacterial genomes, where they might be part of autolysins that need to act on the very same substrate as the archaeal PGH (Fig. 3A). While perhaps not sufficient to confidently identify the bacterial target(s), this stratagem might nonetheless be helpful in whittling down a daunting multitude of potential targets to a priority list of candidate bacteria to test experimentally.

**Figure 3.**
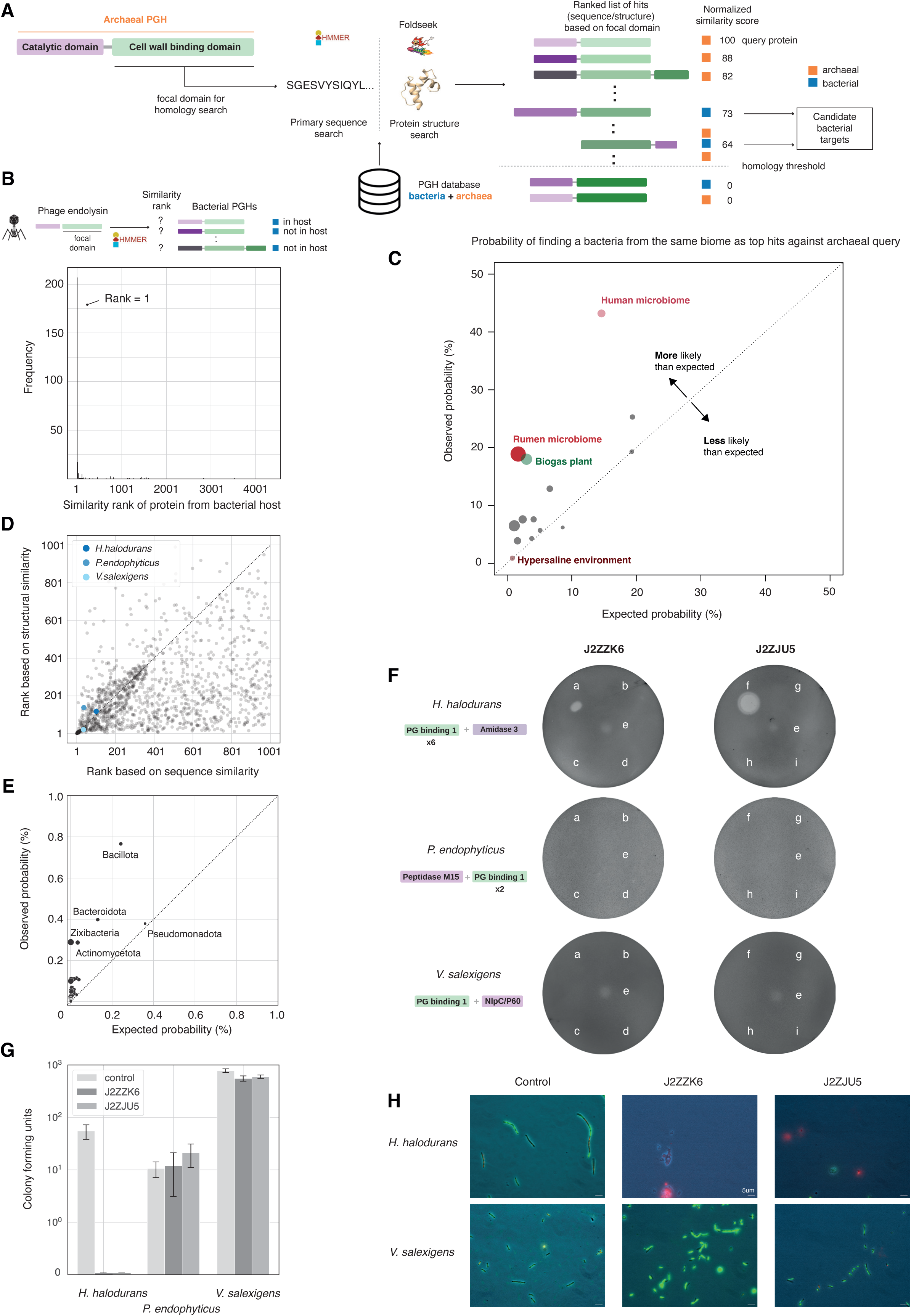
Identification and validation of bacteria targeted by peptidoglycan hydrolases. **A.** Schematic overview of the pipeline for identifying bacteria targeted by peptidoglycan hydrolases. The cell wall/peptidoglycan binding domain or a putative archaeal peptidoglycan hydrolase (PGH) is searched against a database of archaeal and bacterial PGH homologs, using either primary sequence (HMMer) or structural homology (Foldseek) searches. Domain matches are ranked by similarity, normalized to the score obtain by matching against the query protein itself. **B.** The cell wall binding domains of phage endolysins are typically most similar to the cell wall binding domains of their (predicted) bacterial hosts compared to domains from other bacteria. **C.** Probability that, for a given archaeal query protein, bacteria from the same biome are identified amongst the top 10 hits. Biome labels were applied as described in Methods and results are aggregated at the biome level; the size of each circle indicates the likelihood ratio associated with a given biome. **D.** Ranking of homologous bacterial hits against the PG binding 1 domain of J2ZZK6 based on sequence versus structural homology scores (Spearman’s rho = 0.48). **E.** Enrichment of bacterial phyla amongst top ten predicted targets across PGH homologs. See Methods for how enrichment was computed **F.** Results of spotting various supernatants onto lawns of *H. halodurans, P. endophytics,* and *V. salexigens.* a,b: top fraction (>3 kDa) of supernatant from *H. volcanii* expressing an intact (a) or catalytic mutant (b) version of J2ZZK6; c,d: bottom fraction (<3 kDa) of supernatant from *H. volcanii* expressing an intact (c) or catalytic mutant (d) version of J2ZZK6; e: unfiltered supernatant from *Halogranum salarium* B-1; f,g: top fraction (>3 kDa) of supernatant from *H. volcanii* expressing (f) J2ZJU5 or (g) and empty plasmid control; h,i: bottom fraction (<3 kDa) of supernatant from *H. volcanii* expressing (h) J2ZJU5 or (i) and empty plasmid control. **G.** Colony forming assays for different bacteria following treatment with supernatant from *H. volcanii* expressing J2ZZK6, J2ZJU5 or the control (bottom fraction of J2ZZK6-expressing *H. volcanii* H1424 supernatant). **H.** Images of bacterial cells following exposure to *H. volcanii* supernatants expressing J2ZZK6, J2ZJU5 or the control (as above). Bacteria are stained using LIVE/DEAD® BacLight™ bacterial viability staining kit with ingress of red dye into bacterial cells indicative of cells with terminally compromised cell wall integrity.

To explore whether this approach is likely to be informative, we first considered PGHs from phages, where they are known as endolysins. The advantage of phages is that we can predict their bacterial target with reasonable accuracy (or even certainty, in the case of lysogenic phage that have embedded themselves in the genome of their host, where they lie dormant until activated). We used the Virus-Host database (Mihara et al. 2016) to map phages to their hosts, focusing on phages whose host is present in our genome database (Table S3; see Methods). To avoid circularity, we removed all PGHs from the target database that are found in parts of bacterial genomes predicted to be prophages by PhiSpy (Akhter et al. 2012). We then asked how often, when carrying out homology searches with a given phage endolysin, we retrieve a PGH from the bacterial host as the top hit versus a PGH from another bacterium in the same database. Strikingly, the majority of host bacteria appear in the top 10 of the search results (N=256/479, 53%) and the median rank of host bacteria is 1 (Fig. 3B; Table S4). Following the same approach but for the catalytic domain of the phage endolysin also regularly identifies the bacterial host in the top 10, but does so less consistently (N=170/479, 35%; Fig. S2, Table S4).

Encouraged by these results, we proceeded to predict bacterial targets of archaeal PGHs, using both primary sequence and structural homology to assess similarity between bacterial and archaeal PGB domains (Fig. 3A). Below, we focus on our focal archaeal protein (J2ZZK6 from *H. salarium* B-1), but predicted bacterial targets for the entire suite of archaeal PGHs, which are particularly enriched for gram-positive bacteria from the phylum Bacillota (Fig. 3E), can be found in Table S5 (available at https://github.com/srom/archaea-vs-bacteria/blob/main/data/putative_targets.csv.gz).

Considering the top bacterial hits (at both a primary sequence and structural level) for J2ZZK6, we selected three halophilic bacteria for further investigation: *Halalkalibacterium halodurans* (UniProt ID of matched protein: Q9KDB8), *Virgibacillus salexigens* (UniProt: A0A024QGA1), and the moderately halophilic soil dweller *Phycicoccus endophyticus* (UniProt: A0A7G9R4I8, Table S6, Fig. 3D). We focus on these bacteria for three reasons: first, they are commercially available and can been cultured in isolation whereas many of the other top hits are against bacterial genomes assembled from metagenomic data (Table S6); second, their basic growth requirements overlap with those of *H. salarium.* Even though the bacteria do not thrive at salinities that are optimal for *H. salarium –* 2.6-3.4M NaCl (Kim et al. 2011) *–* the archaeon can survive and grow at the lower salinities (up to 1.4M NaCl) preferred by the bacteria. Consequently, these organisms might conceivably encounter each other in their natural niches; third, given that *H. salarium* is adapted to thrive under high salt conditions, it is reasonable to assume that its proteins, including J2ZZK6, are adapted to function in a high-salt environment and might fail to function under low-salt conditions. Testing PGH functionality on bacteria growing in a halophilic medium reduces the risk that media composition scuppers enzyme activity.

To test for PGH-mediated inhibition of putative target bacteria, we cloned J2ZZK6 and expressed it in the model halophilic archaeon *Haloferax volcanii* (strain H1424, see Methods). Supernatant from *H. volcanii* expressing J2ZZK6 did not affect the growth of *P. endophyticus* or *V. salexigens* (Fig. 3F). In contrast, we observed robust effects on the growth of *H. halodurans* (Fig. 3F-H). Importantly, supernatant from *H. volcanii* without the expression plasmid or with a plasmid that carries a mutant version of J2ZZK6 with a compromised catalytic domain (see Methods) did not cause inhibition, suggesting that J2ZZK6 is the cause of, and catalytic activity critical for, inhibition.

Streptococcus zoocin A fatally compromises cell wall integrity and therefore has a bactericidal rather than bacteriostatic effect (Simmonds et al. 1996). To establish whether this is also the case for J2ZZK6, we carried out two complementary tests: colony forming assays and fluorescent live/dead staining. Both tests indicate bactericidal activity against *H. halodurans* (Fig. 3G,H; Fig. S3).

### Bactericidal activity of Halogranum salarium phase supernatant

Is J2ZZK6 secreted by *H. salarium* and does *H. salarium* supernatant inhibit bacterial growth? To address these questions, we spotted filtered *H. salarium* stationary phase (day 6) supernatant onto lawns of *H. halodurans, P. endophyticus,* and *V. salexigens*. *H. salarium* supernatant exhibited mild but consistent bactericidal activity against *H. halodurans* (Fig. 3F). However, bacteria often carry not one but several proteins that can be deployed against competitors. It would therefore be prudent to assume that J2ZZK6 is not the only protein in the *H. salarium* supernatant with bactericidal activity and that the native supernatant assay therefore does not directly implicate J2ZZK6. To investigate whether proteins other than J2ZZK6 might be behind the observed inhibition, we carried out proteomic analysis of supernatants from *H. salarium* cultures in stationary phase (day 6). Using quantitative label-free proteomics, we detect 1133 proteins (Table S7). This set includes J2ZZK6. However, we also detect a second PGH (UniProt ID: J2ZJU5, Fig. 2B), composed of a glucosaminidase domain (PF01832) and two tandem PG binding 1 domains (PF01471) closely related to those in J2ZZK6 (Fig. 2B). Like J2ZZK6, J2ZJU5 carries a signal peptide consistent with active secretion. To test whether J2ZJU5 too exhibits bactericidal activity, we carried out the same suite of experiments described above. We find that J2ZJU5 does indeed kill *H. halodurans* (but not the other tested bacteria) when secreted from *H. volcanii* H1424, and appears more potent than J2ZZK6 under the same conditions (Fig. 3F-H). J2ZJU5 might therefore be partly (or even chiefly) at fault for the bactericidal activity of the native supernatant.

Interestingly, *H. salarium* supernatant also mildly affected the growth of *V. salexigans,* despite the fact that neither J2ZZK6 nor J2ZJU5, heterologously expressed from *H. volcanii*, exhibited bactericidal activity against this target (Fig. 3F). This might suggest that successful targeting of this bacterium by J2ZZK6/J2ZJU5 depends on additional factors not present in the supernatant of the heterologous expression system. Alternatively, *H. salarium* might secrete yet more factors, unrelated to the two PGHs, that compromise *V. salexigans* growth.

### Does the PGB homology approach identify ecologically relevant targets?

The observations above suggest that homology searches against the PGB domain(s) of archaeal PGHs might be useful to identify bacterial targets. Although further experimental work will be required to establish the specificity and sensitivity of this approach, we wondered whether the predicted targets broadly make ecological sense.

To address this question, we made use of the MGnify database, a large compilation of consistently labelled microbiome (meta)genomes (Richardson et al. 2022). We first assigned appropriate biome labels from the MGnify ontology to our original list of PGH-encoding archaea (see Methods). We then searched for bacterial homologs of archaeal PGHs in MGnify, asking whether, for a given archaeal PGH, the top-ranked bacterial hits share the same biome label as the producer more often than expected by chance (see Methods).

For 12 out of 14 biomes, we find evidence for non-random biome correspondence (likelihood ratio > 1, Table S8, Fig. 3C). Interestingly, bacteria from hypersaline environments are not significantly enriched amongst the top targets of archaeal hypersaline PGHs. On closer scrutiny, however, we find that top hits often come from biomes that could be considered frontier environments for halophiles, such as “fermented vegetables”, which – as exemplified by kimchi – can also constitute salt-rich environments. *H. halodurans* itself is assigned the label “Environmental:Aquatic:Marine:Intertidal zone”, the transitional zone where the ocean meets the land between high and low tides. As bacteria tend to increase in relative abundance compared to archaea as salinity drops (Belilla et al. 2021), we propose that PGHs and similar antagonistic tools might be particularly useful in these environments.

## DISCUSSION

We demonstrate above that a diverse assortment of archaea encode peptidoglycan hydrolases; enzymes that target a cellular structure, peptidoglycan, not found in archaea. At least three archaeal PGHs – J2ZZK6 and J2ZJU5 from *H. salarium* B-1, as shown here, and B5ID12 from *A. boonei* (Metcalf et al. 2014) – kill bacteria. As *H. salarium* B-1 was isolated from evaporitic salt crystals from the sea shore of Namhae, Korea, we name the genes encoding J2ZZK6 and J2ZJU5, *woldo* (“moon blade”) and *danwoldo*, respectively, in reference to two historical Korean polearms, the latter having a larger blade.

To identify putative bacterial targets of archaeal PGHs we used a (structural) homology approach, which – pending broader validation – might become a valuable tool for identifying producer-target relationships. Approaches of this kind might be particularly useful for antagonistic interactions, where, for obvious reasons, producer and target are less likely to be found in the same sample, precluding some other common metrics to infer interactions (e.g. recurrent local co-occurrence, as one might expect under syntrophy).

Our findings prompt a series of questions: How are archaeal PGHs (and potentially other archaeal proteins with antibacterial activity) deployed in an ecological context? Woldo (J2ZZK6) and Danwoldo (J2ZJU5) are present in the supernatant of stationary phase *H. salarium* grown in monoculture, suggesting that they are expressed pre-emptively in nutrient-poor conditions, perhaps in anticipation of more competitive times ahead. Similar expression dynamics have previously been observed for halocins – small proteins that are secreted by halophilic archaea to kill other halophilic archaea (O’Connor and Shand 2002). But might expression also be more specifically regulated, i.e. induced by the presence of bacteria? If so, how do archaea sense bacteria and do they sense specific bacteria specifically? Do they express PGHs to defend their niche and compete for resources? Or are bacteria themselves the resource, parcels of nutrition that are unlocked by PGHs?

Whatever the answers, understanding how archaea interact with bacteria – through PGHs and other means – will be key to understanding how archaea persist in and shape polymicrobial ecosystems and to predicting short-and long-term change in the composition of these consortia. Because of their unique metabolic capacities, archaea may often act as keystone species (Moissl-Eichinger et al. 2018), with an outsize influence over community dynamics. Recognizing that archaea can interact in an antagonistic manner with bacteria in their community, actively shaping their niche, will change our appreciation of ecosystem function at equilibrium but also our ability to predict how interventions would play out.

It is worth noting here that we detect PGHs not only in archaea that inhabit niches where bacteria abound, but also in archaea that are usually found in environments where bacteria are scarce. Notably, this includes *H. salarium,* which optimally grows at salt concentrations that are prohibitive for most bacteria (see above). Why would archaea that dominate their niche encode proteins that target bacteria? We speculate that, for these species, PGH might nonetheless be useful when they encounter what, for them, are frontier environments: niches at the edge of their range, where lower salinity allows bacteria to be competitive and archaea survive rather than thrive. In the future, it will be interesting to define the (stress) conditions that trigger PGH expression and how they relate to the ecophysiology of the particular archaeon under investigation.

### A roadmap for the discovery of additional archaeal-bacterial interactions

Our findings suggest, as much by implication as directly, that antagonistic interactions between archaea and bacteria might be pervasive. We think it unlikely that PGHs will turn out to be the sole mediators of archaeal-bacterial antagonism. Bacteria use a large variety of molecular systems to interfere with each other’s growth and survival. The pan-bacterial arsenal includes large, complex molecular machines such as type VI secretion systems, contact dependent inhibition, bacteriocins that can penetrate and depolarize bacterial membranes, and an assortment of small molecules, that we have come to rely on as antibiotics (Smith et al. 2023). Might archaea have access to an equivalent armoury? And how similar is it to that of bacteria?

We suggest four broad strategies to discover additional interactions in the future:

First, peptidoglycan is not the only biological structure that differentiates archaea from bacteria. Other acute points of difference exist, one being the lipids used in archaeal and bacterial membranes: archaeal lipids are made from isoprenoid chains that are linked by ether bonds to a glycerol-1-phosphate backbone. Bacteria, in contrast, predominantly use ester bonds to link fatty acids to a glycerol-3-phosphate backbone (Caforio and Driessen 2017). One might therefore pursue a similar strategy to the one we adopted above and search archaeal genomes for enzymes that specifically interact with bacterial phospholipids.

Second, whereas, at present, we do not know the potentially diverse methods by which archaea kill bacteria, we do know a great deal about how bacteria kill bacteria (Smith et al. 2023). It would be interesting to establish systematically whether systems homologous to known bacterial weaponry are present in archaeal genomes. For a first glimpse into whether archaea might use the same toolkit as bacteria to kill bacteria we searched our archaeal genome collection for homologs of bacteriocins collated in the BAGEL4 database (van Heel et al. 2018). We find that some well-studied bacteriocins, like Lactoccocin 972, are never found in archaea (Fig. 4). Others, like colicin M or sakacin A, are found in archaea in a patchy phylogenetic pattern reminiscent of their distribution in bacteria (Fig. 4, Table S9) (Azevedo et al. 2015; Sharp et al. 2017) and might therefore be good candidates for archaea-produced bacteriocins. However, candidate status here is less cogent than it is for PGHs and needs to be interpreted with care. Importantly, the link to bacteria is less specific: proteins as well as small metabolites could be active against bacteria but evolved to fight against other archaea, or for some other purpose altogether. To what extent these bacteriocin homologs have evolved in and are used by archaea to target bacteria will have to be established experimentally. In this context, it is worth pointing out a recent discovery of predicted pore-forming toxins (along with murein transglycosylases) in metagenome assembled genomes (MAGs) from the phylum Woesearchaeota, which also points to antagonistic interactions with bacteria (Vigneron et al. 2022).

**Figure 4.**
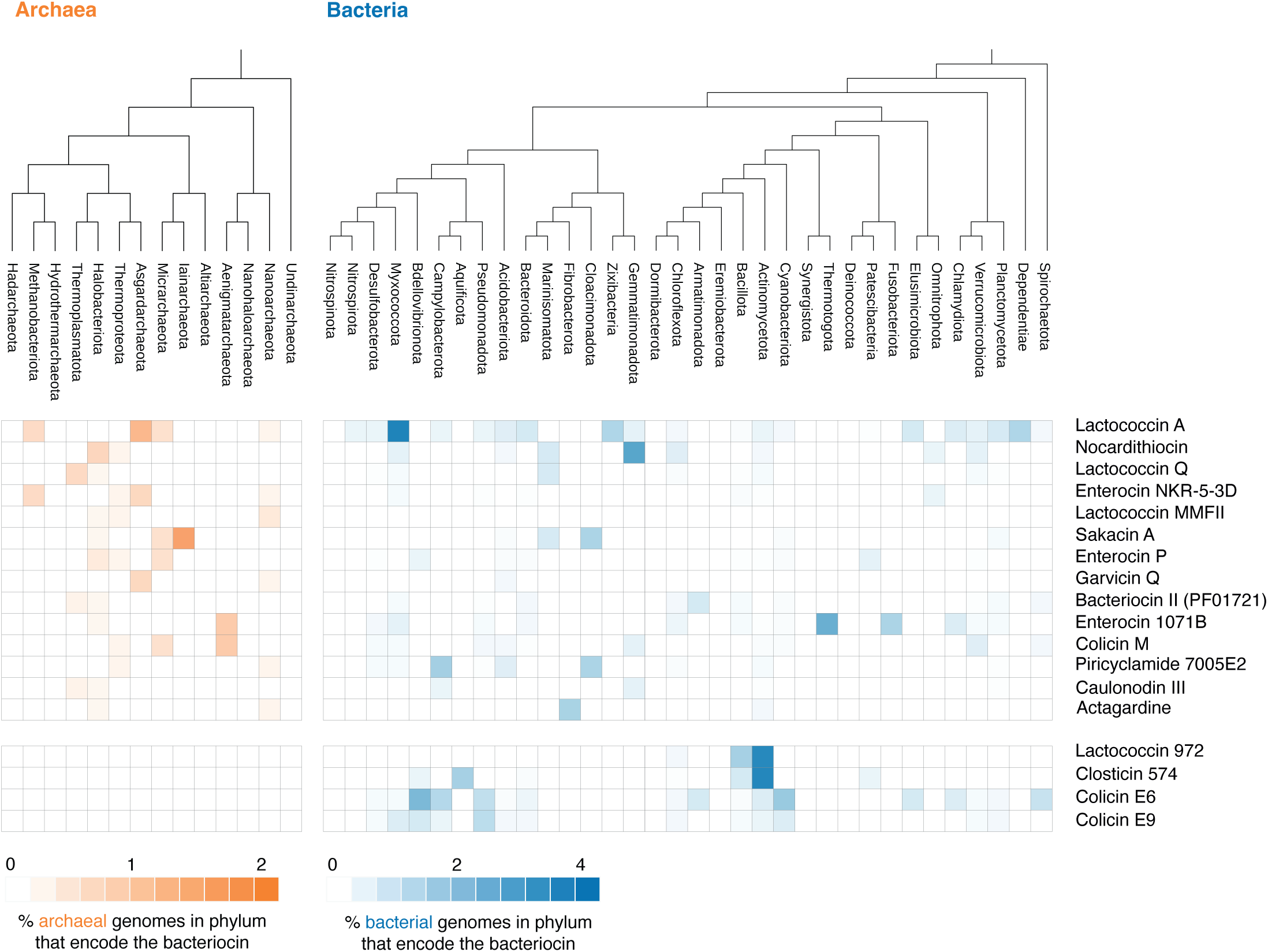
Distribution of bacteriocin/archaeocin homologs across bacteria and archaea. The relative abundance of different bacteriocin/archaeocin homologs is shown aggregated at the phylum level.

Third, the two strategies above rely on homology to known bacterial systems as a starting point. These approaches, by themselves, will not reveal novel systems, especially systems unique to archaea. They may, however, provide critical seeds for discovery. A pertinent example here is the recent avalanche-like progress made in identifying systems involved in conflicts between phage and bacteria: Genes involved in phage defense often cluster in bacterial genomes. Genes of unknown function that repeatedly co-occur with known defense genes were therefore predicted to be involved in anti-phage defense themselves – a guilt-by-association hypothesis that led to the discovery of multiple new phage defense systems (Doron et al. 2018). As yet, we have too few landmarks (archaeal proteins with antibacterial function) to carry out similar analyses. As the number of landmarks grow, this might become a fruitful avenue for investigation. Provided, of course, that genes involved in antibacterial activity also have a tendency to cluster, which is by no means guaranteed.

Finally, bottom-up experimental approaches, including co-culture experiments, will remain invaluable to enable discovery of novel, uniquely archaeal proteins and small molecules with antibacterial activity, to provide the foundations for guilt-by-association approaches, to understand how and to what effect these tools are deployed against bacterial targets, and to assess whether novel archaeal antibacterials might be promising leads for future clinical applications.

## METHODS

### Prokaryotic genome database

We assembled a database of 54,346 prokaryotic genomes (50,640 bacteria and 3,706 archaea), constructed as a phylogenetically balanced subset of release 214 of GTDB, the Genome Taxonomy Database (Parks et al. 2021), selecting a single representative strain per species (as per the representative strain flag available in GTDB) and a maximum of ten species per genus, prioritizing those species with most strains. A full list of genomes and corresponding assemblies is provided in Table S1.

### Identifying peptidoglycan hydrolases

We define peptidoglycan hydrolases as proteins made of at least one Pfam catalytic domain known to cleave peptidoglycan, and at least one Pfam cell wall binding domain, cross-referencing our manually curated list with literature on bacterial autolysins (Mitchell et al. 2021). Manual curation was based on a domain’s Pfam description: all domains mentioning peptidoglycan hydrolase activity or peptidoglycan binding activity have been included. A full list of qualifying Pfam domains is available as Table S10. Proteins matching this definition were found by systematically searching our database of prokaryotic genomes using Pfam’s Hidden Markov Models (HMMs) with the HMMER software suite (hmmer.org). Signal peptides were predicted with SignalP 6 (Teufel et al. 2022). The full list of proteins found is available as Table S2.

PGHs were assigned to main chromosome, plasmid, contig, etc. as follows. For complete genomes, existing annotations were used to assign the PGH to a chromosomal or plasmid location. For incomplete genomes, the size of the contig was compared to the median contig size of the assembly. Contigs below the median size were considered “small contigs”. Scaffold level genomes were excluded from this analysis.

### Phylogenetic trees

Cladograms (Fig. 1C, Fig. 4) are derived from the bacterial and archaeal trees from GTDB: all named phyla were kept (i.e., excluding automatically generated phyla name composed of numbers and letters). Phyla with the same name but differentiated by a letter (such as Pseudomonata, Pseudomonata_A, Pseudomonata_B, etc.) were grouped into a single phylum (e.g. Pseudomonata).

Protein domain trees (Fig. 2) were constructed by selecting the top 1,000 hits to the peptidase M23 and PG binding 1 domains of protein J2ZZK6, respectively. Sequences were aligned with MAFFT using the L-INS-I strategy (Katoh and Standley 2013). Alignment positions with less than 35% aligned residues were trimmed with trimAl (Capella-Gutiérrez et al. 2009). Maximum likelihood trees were built with IQ-TREE 2 (Minh et al. 2020) with automatic model finder (“-m MFP” option) and UFBoot2 approximate bootstrap values (“-B 1000” option). Only the bacterial clades branching with archaea plus one outgroup were kept for clarity.

The tree in Fig. S1 is a subset of the GTDB archaeal tree containing only Halobacteriota species present in our database. All trees were rendered with iTOL (Letunic and Bork 2024).

### Phage-Host database

Bacteriophages and their host were retrieved from the Virus-Host database (Mihara et al. 2016). Among these bacteriophages, only the ones whose host belongs to our genome database were kept, using the bacterium’s NCBI taxonomy ID to perform the matching (Table S3). We then considered phage genomes that contained at least one endolysin.

### Predicting bacterial targets of archaeal PGHs and phage endolysins

To identify the likely bacterial targets of phage or archaeal PGHs, we used HMMER (phmmer) to carry out homology searches against our prokaryotic database, using only the cell wall binding domain as a query and keeping proteins with an E-value ≤ 1e-6. Following prediction of tertiary structures of the full-length homologous target proteins using ColabFold (Mirdita et al. 2022), target proteins were ranked either by primary sequence or structural homology to the archaeal query domain using the HMMER and Foldseek output (Kempen et al. 2024), respectively (Fig 3D).

### Biome predictions

To evaluate how often our target search approach identifies bacteria that inhabit the same or a similar environment to the archaeal PGH producer, suggestive of ecologically relevant cross-domain interactions, we require a set of consistent biome labels across bacterial and archaeal genomes. Controlled labelling of this type has been done for genomes in the MGnify database (Richardson et al. 2022). We matched proteins from this dataset to the PGH proteins in our database using phmmer with an e-value threshold of 1e-6, in effect importing labels from MGnify into our database using homology as a guide. Many of the Mgnify biome labels are broad (e.g. Environmental:Aquatic:Marine). Therefore, we picked the three top biome labels per PGH protein. For each archaeal PGH protein, we extracted the biome labels of the top ten predicted bacterial targets and asked whether the probability of the archaeal producer label to appear in the top ten bacterial targets is statistically different from randomly sampling ten labels from the entire bacterial PGH dataset.

### Phylum enrichment

To establish whether particular phyla were enriched amongst predicted bacterial PGH targets, we considered the entire set of bacterial PGHs in our database (Table S2) and computed the probability of retrieving a hit from a particular phylum by chance and multiplied this number by ten to generate an expected probability of a protein to appear amongst the top 10 hits. We then considered the bacterial PGH hits for each archaeal PGH (Table S5) and computed the observed probability of a phylum to appear in the top 10 (see https://github.com/srom/archaea-vs-bacteria/blob/main/notebook/figureS2.ipynb for further details).

### Antimicrobial proteins shared between archaea and bacteria

To identify putative antibacterial proteins in archaea we searched in our database for homologs to antimicrobial proteins present in the BAGEL4 database (van Heel et al. 2018). Proteins in BAGEL4 were clustered using CD-HIT (Fu et al. 2012) to group together proteins identical or similar in sequence (90% similarity; option -c 0.9) and this consolidated set of proteins were used as queries to search against our database using HMMER, with an E-value threshold of 1e-6 (Table S9). The heatmap in Fig. 4 only includes proteins present in both archaea & bacteria in at least 2 genomes and 2 phyla in each domain, plus four selected bacteriocins not present in archaea. The set shared by archaea and bacteria was further reduced to exclude the following proteins labeled as bacteriocins in BAGEL4: peptidoglycan hydrolases; Linocin M18, which has known non-bactericidal functions in *Pyrococcus furiosus* (Namba et al. 2005); FlvA2f and FlvA2h, which have been shown to have no bactericidal activity (Zhao and van der Donk 2016); and comX homologs, pheromones involved in quorum sensing (Schneider et al. 2002), since no bactericidal activity has been described.

### Strains and culture conditions

*Halogranum salarium* B-1 (DSM-23171), *Halalkalibacterium halodurans* (DSM-18197), *Phycicoccus endophyticus* (DSM-100020), and *Virgibacillus salexigens* (DSM-11483) were obtained from the Deutsche Sammlung von Mikroorganismen und Zellkulturen (DSMZ). *Haloferax volcanii* H1424 was a gift from Thorsten Allers (University of Nottingham). *H. salarium* B-1 was grown in DSMZ media recipe 1377 (13% w/v NaCl). *H. volcanii* was grown in medium Hv-YPC 18% w/v NaCl as previously described (Dattani et al. 2022). The three bacteria *H. halodurans*, *P. endophyticus* and *V. salexigens* were grown in Hv-YPC 8% w/v NaCl. Note here that *Halogranum salarium* B-1 can also grow, albeit not optimally, at 8% w/v NaCl. Pre-cultures from −80°C glycerol stocks were grown in 3 mL volume in 15 mL Falcon™ tubes. Cultures were grown in 20 mL volume in 100 mL Erlenmeyer flasks. *H. salarium B-1*, *H. halodurans*, *P. endophyticus* and *V. salexigens* were grown at 37°C, while *H. volcanii* was grown at 45 °C, all in a shaking incubator at 180 rpm.

### Transformation of Haloferax volcanii

DNA sequences corresponding to J2ZZK6 and J2ZJU5 were amplified from *H. salarium* B-1 genomic DNA by PCR (see Table S11 for primers) and cloned into *H. volcanii* plasmid pTA1392 (a gift from Thorsten Allers) as follows: the plasmid was digested with NdeI and BamHI and subsequently dephosphorylated using the NEB shrimp alkaline phosphatase to prevent vector circularisation after transformation. The digested PCR product and the dephosphorylated vector were ligated using T4 DNA ligase at a 3:1 molar ratio. The ligation mixture was electroporated into *E. coli DH5a*, and transformed clones were selected on LB agar + 100 ug/mL carbenicillin. The resulting plasmids were extracted using the NEB miniprep plasmid extraction tool and transformed into *H. volcanii* H1424 by following an established transformation protocol (Dattani et al. 2022). To serve as a control, the original pTA1392 without a gene insert was similarly transformed into *H. volcanii* H1424 using the same protocol. Finally, we sought to construct a catalytic mutant of J2ZZK6. Introducing a point mutation via PCR proved difficult because the high GC content of the gene encoding J2ZZK6 prevented design of workable primers. Instead, we synthetised a suitable DNA fragment containing the necessary mutation and attempted to introduce the fragment into pTA1392 via Gibson assembly. None of the mutant we screened had a successful integration event. However, we fortuitously isolated a mutant plasmid with a 177 bp deletion which covered the catalytic centre of J2ZZK6. All plasmids were sequenced (via Plasmidsaurus) prior to deployment to confirm expected integration events.

### Inhibition assays

*H. salarium* cultures were inoculated at 0.01 OD600 from overnight cultures, into 20 mL 13% w/v NaCl media in 100 mL Erlenmeyer flasks and grown for 6 days. *H. volcanii* cultures were inoculated at 0.01 OD600 from overnight cultures, into 20 mL Hv-YPC 13% w/v NaCl + 3 mM Tryptophan to induce expression, in 100 mL Erlenmeyer flasks overnight. Supernatant was extracted by centrifugation of liquid cultures of *H. salarium* or *H. volcanii* at 4,000g for 10 minutes. Supernatant was subsequently passed through 0.2 µm Minisart PES sterile filters to remove cells. Supernatant was then concentrated using 3,000 MWCO PES membranes (Vivaspin 20) by centrifugation for 45 minutes at 4,000g. Volume in the top compartment was reduced from 20 mL to 1 mL (20x) and is enriched in molecules with a molecular weight >3 kDa. The bottom compartment holds the flow through. Spotting assays on target lawns were carried as previously (Shand 2006). Briefly, target bacterial cells were incorporated into a soft agar top layer, which was poured hot into a plate containing hard agar of the bacterial medium. Supernatant was spotted on top of the plate straight after the top layer had solidified. Plates were left for colonies to grow at 37°C and imaged after 24h and inspected for clearings.

### Colony-forming units

Bacterial cultures of *H. halodurans*, *P. endophyticus* and *V. salexigens* were grown as described above to an OD of 0.5. Concentrated supernatant containing proteins J2ZZK6 and J2ZJU5, respectively, were collected as described above. The flow through from supernatant concentration of J2ZZK6 was used as control. A volume of 1 µL of bacterial culture was added to 9 µL of supernatant in Starlab PCR tubes, for a total of 18 tubes (3 bacteria, 3 conditions and 2 replicates). The mixtures were incubated for 1 hour at 37°C in a shaking incubator at 180 rpm. Two 1:100 serial dilutions were performed in bacterial growth media bringing the final dilution factor to 10,000. A volume of 100 µL was plated on pre-warmed solidified bacterial media. Plates were incubated for 36 hours at 37°C and imaged and colonies counted using ImageJ (protocol: <u>10.17504/protocols.io.f2mbqc6</u>).

### Imaging and live/dead staining

Bacterial cultures of *H. halodurans* and *V. salexigens* were grown as described above to an OD of 0.5. 1 mL of culture was centrifuged for 10 minutes at 4,000g. The pellet was resuspended in 500 µL of culture media. Concentrated supernatant from cultures expressing proteins J2ZZK6 and J2ZJU5, respectively, were collected as described above. The flow through from supernatant concentration of J2ZZK6 was used as control. A volume of 1 µL of bacterial culture was added to 9 µL of supernatant in Starlab PCR tubes, for a total of 6 tubes (2 bacteria, 6 conditions). The mixtures were incubated for 1 hour at 37°C in a shaking incubator at 180 rpm. Dyes and protocols from LIVE/DEAD BacLight bacterial viability kits were used. Briefly, equal volumes of SYTO 9 dye (3.34 mM) and Propidium iodide (20 mM) were mixed. The mixture was diluted 1:10 in deionised water. A volume of 0.3 µL was added to each of the 9 tubes and incubated in the dark at room temperature for 15 minutes. A volume of 3 µL of the stained bacterial suspension was trapped between a microscopy slide and a coverslip. Observations were carried out using a Leica DMRB upright microscope with a Leica 100x/1.30 Oil PL Fluotar objective. We used two Semrock Penta florescence filters (FITC 485/20, TRITC 560/25). Acquisition was performed with the Hamamatsu Orca camera using software MicroManager (Edelstein et al. 2010) with 10 ms exposure for the fluorescence channels and 25 ms for the brightfield channel. Composite images were assembled with ImageJ.

### Liquid chromatography–tandem mass spectrometry (LC-MS) sample preparation

Supernatant was extracted from 20 mL cultures of *H. salarium* grown in 100 mL Erlenmeyer flasks for 6 days, in biological triplicates. Secreted proteins were purified following chloroform methanol extraction in the presence of protease inhibitor (cOmplete protease inhibitor cocktail from Sigma). For ease of manipulation, several 200 µL aliquots were processed in parallel in 2 mL Eppendorf tubes. Each aliquot was mixed with 800 µL methanol, 200 µL chloroform, 600 µL of purified water and vortexed until solution turns white. Precipitated proteins were centrifugated at 17,000g for 10 minutes. Next, aqueous supernatant was removed while being careful not to disturb the white protein pellet. Finally, 860 µL of methanol were added and the solution mixed gently. Following 5 min of centrifugation at 17,000g for 5 minutes the chloroform methanol mix was discarded and the pellet allowed to dry for up to 5 minutes. 100 µg of the proteins obtained were processed into purified peptides using the PreOmics iST kit, following manufacturer’s instruction including a 1.5 h digestion step. Purified peptides were stored in LOAD buffer at −80°C until injection.

### LC-MS sample processing & acquisition

Samples were injected and data acquired in single replicate injections as follows: Chromatographic separation was performed using an Ultimate 3000 RSLC nano liquid chromatography system (Thermo Scientific) coupled to an Orbitrap Exploris 240 mass spectrometer (Thermo Scientific) via an EASY-Spray source. Peptide solutions were injected directly onto the analytical column (Self packed column, CSH C18 1.7µm beads, 150μm × 35cm) at working flow rate of 1.3μL/min for 8 minutes. Peptides were then separated using a 69-minute stepped gradient: 0-25% of buffer B for 49 minutes, 25-42% of buffer B for 20 minutes (composition of buffer A – 95/5%: H_2_O/DMSO + 0.1% FA, buffer B – 75/20/5% MeCN/H_2_O/DMSO + 0.1% FA), followed by column conditioning and equilibration. Eluted peptides were analysed by the mass spectrometer in positive polarity using a data-independent acquisition mode as follows: an initial MS1 scan was carried out at 120,000 resolution for 25ms in profile mode, m/z range: 409.5-1650 and normalised AGC target: 300%. This was followed by sequential MS2 acquisition and fragmentation of ions at 30,000 resolutions in centroid mode over 30 variable windows, m/z range: 145-1450, normalised AGC target: 2000% and normalised collision energy: 27%. Total run acquisition time was 84 minutes.

### LC-MS data processing

Data were processed using the Spectronaut software platform (Biognosys, v17.7.230531) (Bruderer et al. 2015). Analysis was carried out in direct DIA mode as follows:

1. Pulsar Search: library generation and database searching were carried out using default settings for a tryspin/p specific digest as follows and with the following adjustments - missed cleavage rate set to 3 and variable modifications allowed for methionine oxidation, protein N-terminal acetylation, asparagine deamidation and cyclisation of glutamine to pyro-glutamate. PSM, Peptide and Protein group FDR = 0.01. Searches were run against a user generated protein sequence database for *H. salarium* B-1 (RefSeq: GCF_000283335.1).
2. *DIA analysis*: a mutated decoy database approach was employed with protein q-value cut-off for the experiment set to 0.05 at the identification level. Quantification set to MS2 with proteotypicity filter set to only proteotypic peptides with no value imputation strategy employed. Protein quantification method set to MaxLFQ (Cox et al. 2014).

## CODE & DATA AVAILABILITY

Code and supporting datasets can be found at https://github.com/srom/archaea-vs-bacteria.

### ACKNOWLEDGEMENTS

We thank Thorsten Allers for *H. volcanii* strains and plasmids, Charazad Taissir for help and advice, and the MRC LMS Proteomics facility for carrying out mass spectrometry experiments. Work in the Warnecke lab is made possible by core funding from the UKRI Medical Research Council (MC-A658-5TY40).

## SUPPLEMENTARY TABLES

**Table S1.** Genomes in the phylogenetically balanced database, including 3,706 archaeal and 50,640 bacterial genomes.

**Table S2.** Peptidoglycan hydrolases detected in bacteria and archaea.

**Table S3.** Phage-bacterial host assignments from on Virus-Host database for the bacterial genomes in our database.

**Table S4.** Correspondence between phage endolysin domains and homologous domains in the bacterial host.

**Table S5.** Predicted bacterial targets based on sequence homology for all archaeal peptidoglycan hydrolases. This table is available at https://github.com/srom/archaea-vs-bacteria/blob/main/data/putative_targets.csv.gz.

**Table S6.** Domains homologous to J2ZZK6 and J2ZJU5 amongst archaea/bacteria in the database.

**Table S7.** Proteins identified in stationary phase (day 6) supernatant of *H. salarium* B-1 cultures.

**Table S8.** Enrichment of MGnify biome labels between archaeal queries and their putative bacterial targets.

**Table S9.** Archaeal homologs of bacteriocins.

**Table S10.** List of domains belonging to putative peptidoglycan hydrolases. Proteins containing at least one catalytic domain and at least one cell wall binding domain from this list were used for homology searches in archaeal genomes.

**Table S11.** Primers used for the amplification of J2ZZK6 and J2ZJU5

---

**Figure S1.**
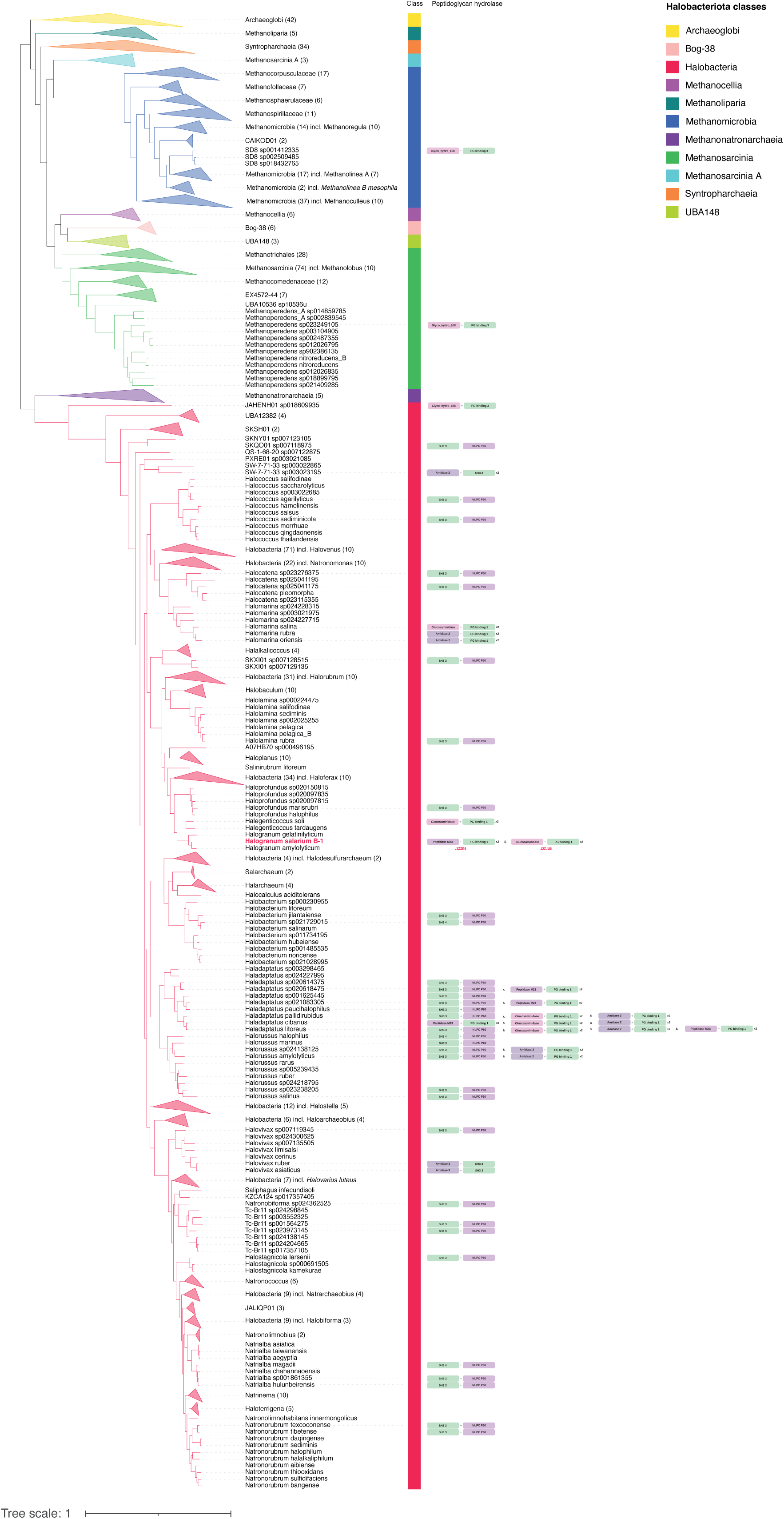
Presence/absence of peptidoglycan hydrolase homologs across the phylum Halobacteriota. The tree shown is based on the GTDB archaeal tree but pruned to only contain Halobacteriota species present in our database (Table S1).

**Figure S2.**
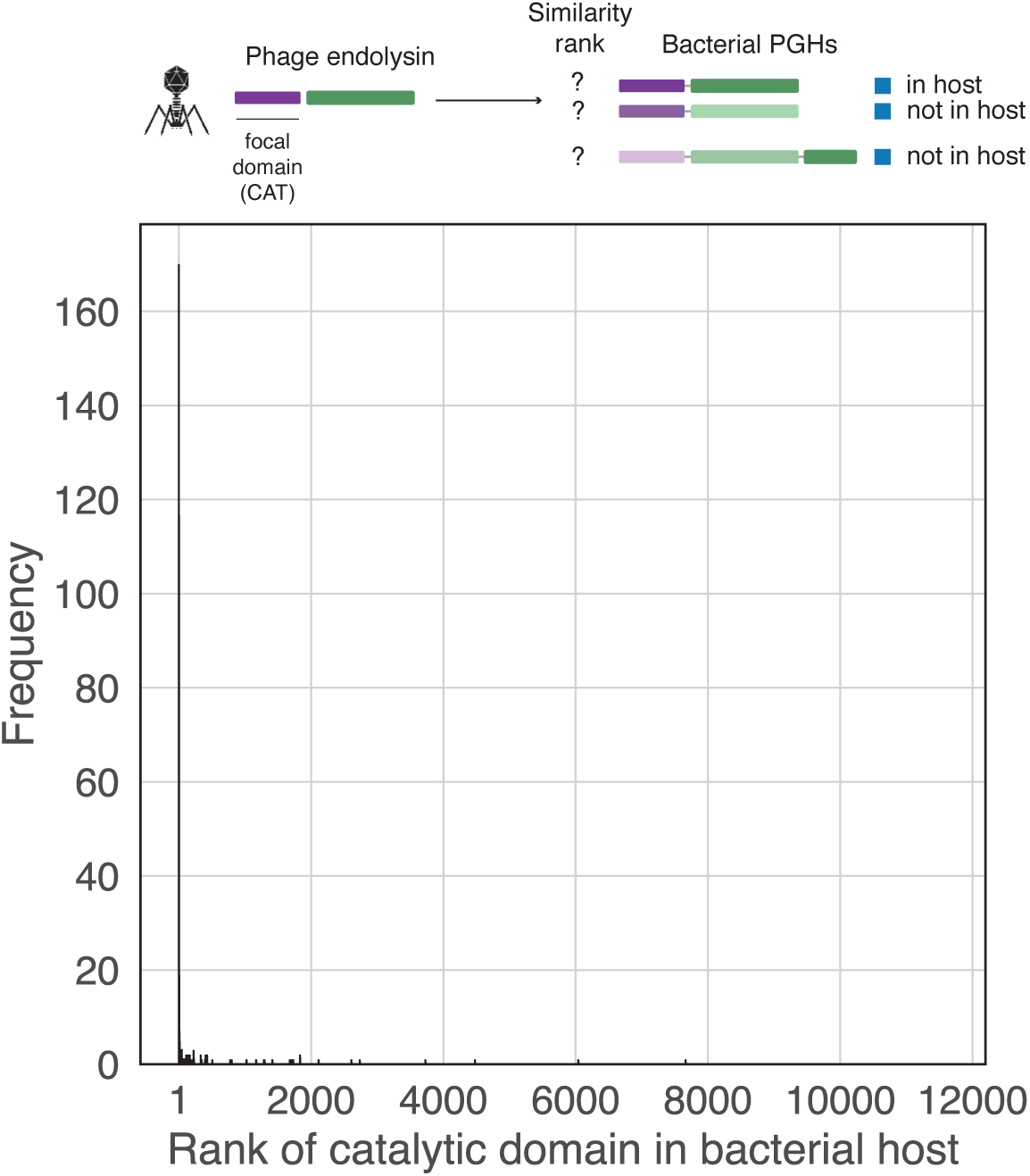
Rank of the bacterial host peptidoglycan hydrolase catalytic domain when searching our bacterial database of with the catalytic domain of a given phage endolysin. This figure corresponds to Fig. 3B but for the catalytic (CAT) instead of the cell wall binding domain.

**Figure S3.**
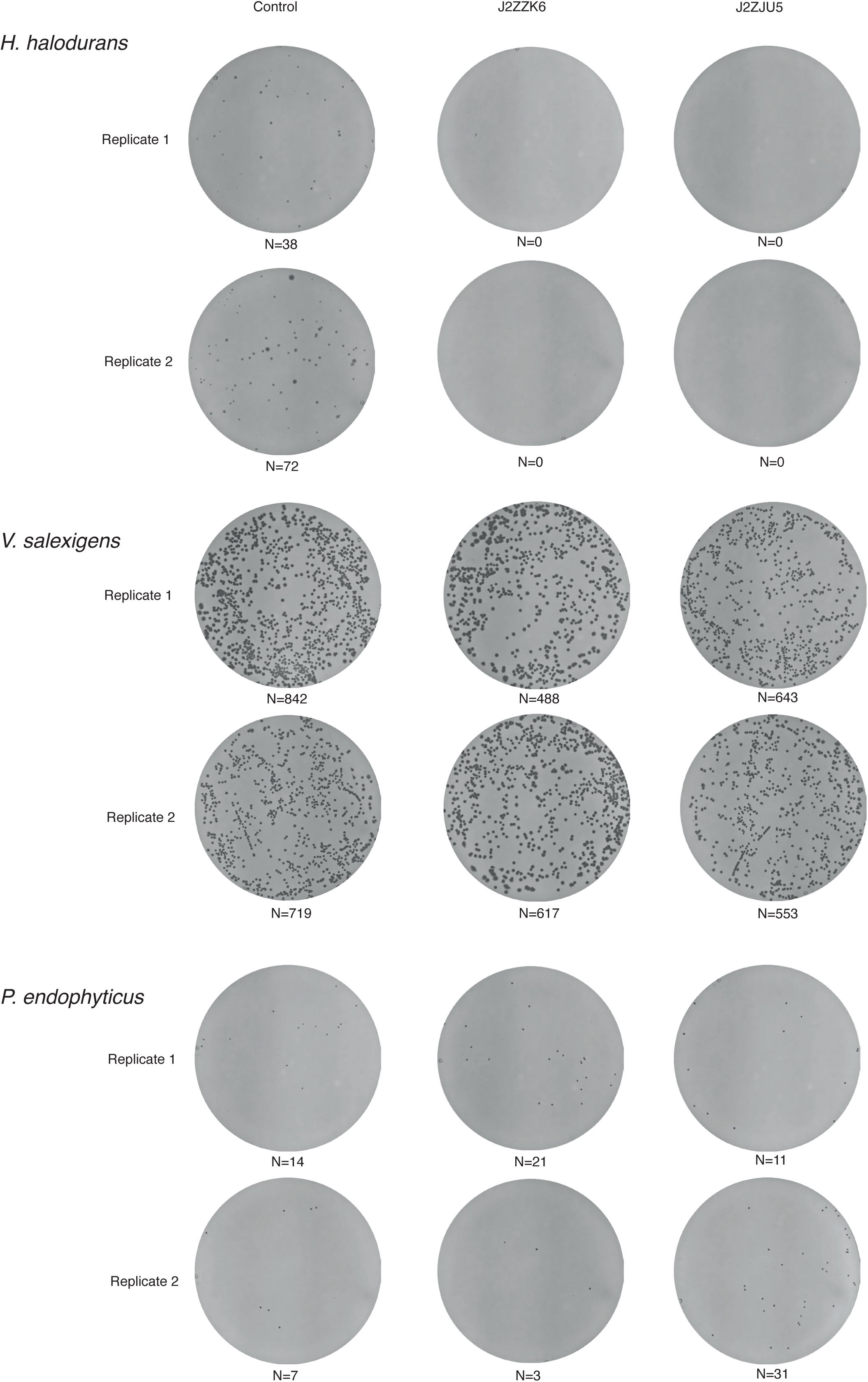
Plates used to determine colony forming units, following exposure to supernatant containing J2ZZK6 or J2ZJU5.

